# The essential genome of the crenarchaeal model Sulfolobus islandicus

**DOI:** 10.1101/408351

**Authors:** Changyi Zhang, Alex P. R. Phillips, Rebecca L. Wipfler, Gary J. Olsen, Rachel J. Whitaker

## Abstract

*Sulfolobus islandicus* is a model experimental system in the TACK superphylum of the Archaea, a key lineage in the evolutionary history of cell biology. Here we report a genome-wide identification of the repertoire of genes essential to *S. islandicus* growth in culture. We confirm previous targeted gene knockouts, uncover the non-essentiality of functions assumed to be essential to the *Sulfolobus* cell, including the proteinaceous S-layer, and highlight key essential genes whose functions are yet to be determined. Phyletic distributions illustrate the potential transitions that have occurred during the evolution of this contemporary archaeal cell and highlight the sets of genes that may have been associated with each transition. We use this comparative context as a lens to focus future research on archaea-specific uncharacterized essential genes for which future functional data would provide valuable insights into the evolutionary history of the contemporary cell.

---

41 years ago, Woese and Fox identified the Archaea as a novel microbial lineage distinct from Bacteria^1^. The same year, Woese and Fox proposed a model of cellular evolution in which early cellular life diverged in two directions, one to the Bacteria and the other to LEACA, the <u>L</u>ast <u>E</u>ukaryotic and <u>A</u>rchaeal <u>C</u>ommon <u>A</u>ncestor, which was subsequently split to form the Archaea and Eukaryota domains^2,3,4^. Increases in genome and metagenome sequence data continue to refine this picture, providing reinforcement for many of its key aspects, improving phylogenetic sampling, and providing additional details^5-12^. The tree of life itself has evolved with the addition of new lineages whose gene content and phylogenetic reconstruction suggests that the <u>T</u>haumarcheota, <u>A</u>igarchaeota, <u>C</u>renarchaeota, and <u>K</u>orarchaeota (TACK) lineage of Archaea may hold the esteemed position of sharing a more recent common ancestor with the Eukaryota domain than other archaeal groups^5,6,13–16^.

Today the tree of life provides a framework for studying the evolution of cellular complexity. Genomics and metagenomics provide data on the distribution of genes across this tree and in doing so provide an understanding of the origins and evolutionary dynamics of gene sequences. However, phyletic distributions fall short of establishing the functional evolutionary history of the cell since gene presence does not link directly to function. Truly mapping evolution of today’s complex contemporary cells involves a comparative approach in which functional cellular systems and the interactions of their constituent components are examined at a molecular level in organisms representing key evolutionary lineages across the tree of life.

As a step in that direction, we take a genome-wide functional approach to define 441 genes essential to the growth of *S. islandicus*. *Sulfolobus*, a thermoacidophilic genus from geothermal hot springs, is one of the few organisms within the TACK archaea that can be cultured and is genetically tractable, and it is the most developed model for studying the biology of cells in this lineage. We highlight surprises revealed by subsequently examining the function of essential and non-essential genes in this model organism, including the non-essentiality of the S-layer protein found to be present in most cells in the archaeal domain^17^. As a step toward comparative functional cell biology, we illustrate the stages of evolution of the essential gene repertoire of the contemporary functional archaeal cell and provide a lens through which to focus attention on uncharacterized genes that will enable further characterization of transitions in cellular evolution.

## Results and Discussion

### Identifying essential genes in the genome of *S. islandicus* through Tn-Seq

We established three independent genome-wide disruption libraries in an agmatine-auxotrophic strain of *S. islandicus* M.16.4 by using a modified *in vitro* transposon mutagenesis system derived from Tn5 (Epicentre, USA). The transposable element was comprised of a nutritional marker cassette, *SsoargD* (arginine decarboxylase derived from *Sulfolobus solfataricus* P2), flanked by two 19-bp inverted repeats (Fig. 1a). After electroporation-mediated transformation of ArgD^-^ cells with the EZ-Tn5 transposome, cells were allowed 10 days of growth on rich media. While valuable information about metabolic and regulatory genes could have been gained by comparing results from different media conditions, we restricted this study to one rich medium to focus on central cellular rather than metabolic functions. Insertion locations were determined via genome tagging and fragmentation (“tagmentation”) on colony pools, followed by amplification and sequencing of the junction sites, which were then mapped onto the genome. In all, 89,758 unique insertion events with at least 3 reads each were identified across all three libraries, corresponding to an average of one insertion every 29 base pairs and an average expected 29 insertions in each annotated protein-coding gene (see Methods; Supplementary Table 1 contains colony, insertion, and read counts for each library while all insertion locations can be found in Supplementary Dataset 1).

**Figure 1:**
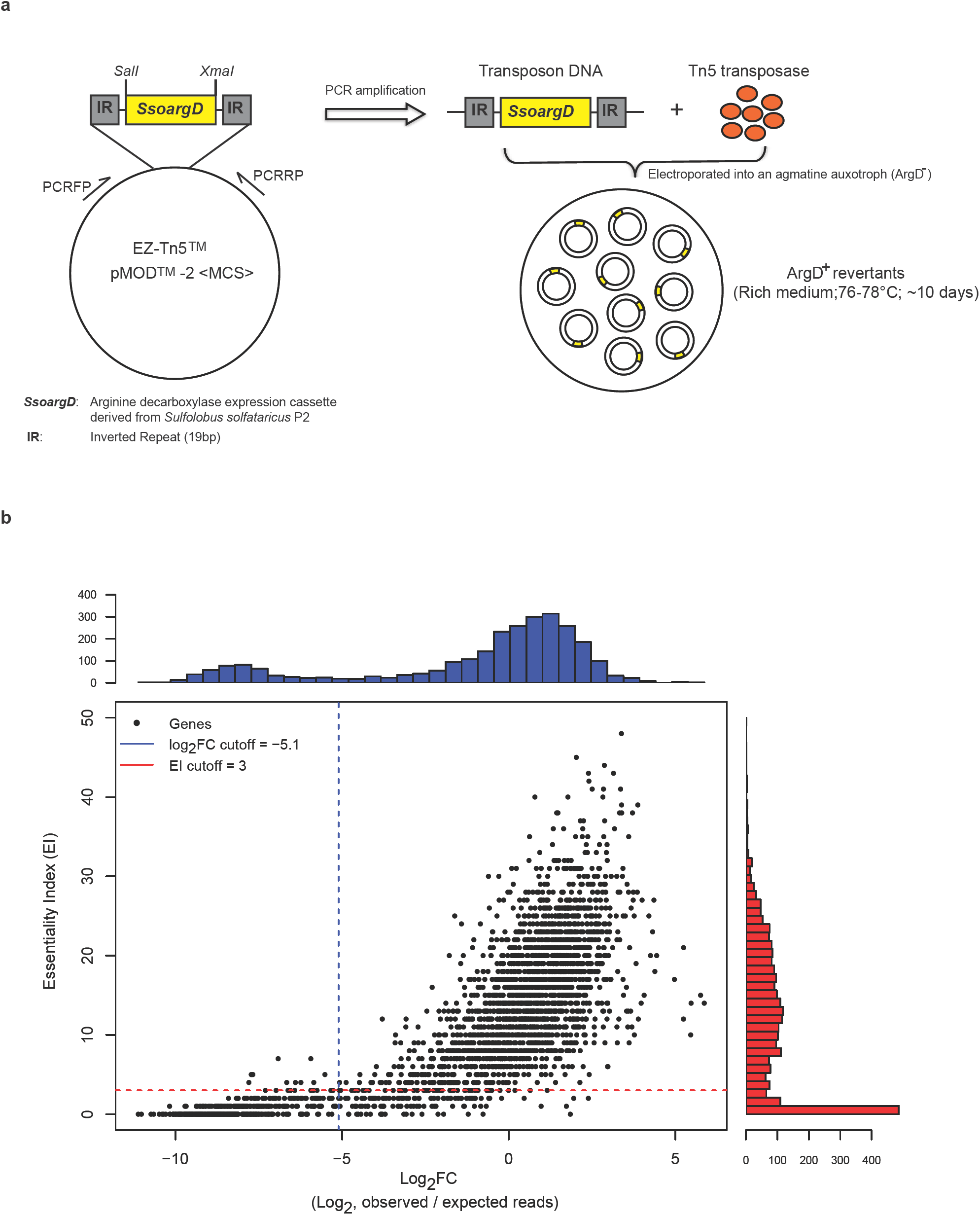
Defining the essential genes in *S. islandicus* M.16.4. **a,** Schematic overview of the genome-wide transposon mutagenesis strategy. **b,** Evaluation of gene essentiality by two computational programs: ESSENTIALS^18^ and Tn-Seq-Explorer^19^. Points indicate individual genes plotted according to the scores returned by each program. Histograms indicate the number of genes of a particular score, and the dotted lines indicate the recommended cutoffs returned by each program as the local minimum between the essential and non-essential score distributions. Essential genes meet both criteria (lower-left quadrant). The protein-coding genes that only met the ESSENTIALS or Tn-Seq-Explorer criteria were deemed as “unas- signed candidates” leaving the rest as likely non-essential to *S. islandicus* M.16.4 growth under these conditions. A complete list of the log2FC and EI for the *S. islandicus* M.16.4 genes from the combined mutant libraries are provided in Supplemental Dataset 2.

Essential genes were predicted to be significantly underrepresented in the insertion locations extracted from the transposon mutagenesis and sequencing data (Tn-seq). It is important to note that this may make them indistinguishable from genes that are not strictly essential for growth, but instead cause a severe growth defect, and thus our definition of “essential” extends to these genes too. To determine the statistical separation between essential and non-essential genes, we used a combination of two programs: ESSENTIALS^18^ and Tn-Seq Explorer^19^. Both methods report essential gene candidates by separating essential and non-essential genes into a bimodal distribution of scores. ESSENTIALS does so by calculating a log ratio of observed and expected reads in each gene (log_2_FC), while Tn-Seq Explorer uses a sliding window approach to examine the absolute number of insertions in and around genes and calculates an Essentiality Index (EI) for each. The former tends to underestimate the number of essential genes, while the latter tends to overestimate^19^. 445 genes lie within the suggested range for both methods (log_2_FC ≤-5.1 and EI<4), leaving 178 genes within only one range, or “unassigned” as essential or non-essential. The remaining 2,105 protein-coding genes are likely non-essential for growth under these conditions (Fig. 1b and Supplementary Dataset 2). Three genes identified as essential through automated methods were additionally removed because misplaced multiply mapped reads falsely reduced read count (*M164_0862*, *M164_1012*, and *M164_1867*; see Supplementary Table 2). Assignments of all genes to categories with their scores for each method are listed in Supplementary Dataset 2.

### Genetic confirmation of essential gene criteria

To support our informatic essentiality/non-essentiality criteria, 129 genes were compared with gene knockout studies performed in our model *S. islandicus* M.16.4 and another two genetically tractable *S. islandicus* strains: RYE15A and LAL 14/1 (Supplementary Table 3). We were unable to acquire knockouts for 42 of 45 predicted essential genes in this set. Two exceptions, *topR2* (*M164_1245*) and *apt* (*M164_0158*), were identified to have significant growth defects on plates once they were knocked out (Supplementary Fig. 1c, 2a and ^20^), likely resulting in their under-representation in our transposon library. The third, *cdvB3* (*M164_1510*), a paralog of *cdvB*, may be incorrectly called essential in our Tn-seq analysis. We can readily obtain *cdvB3* disruption mutants (Supplementary Fig. 3b) and the growth of a *cdvB3* mutant strain is indistinguishable from the wild-type strain (data not shown), thus this gene was removed from the essential gene list. An explanation of why this gene is mischaracterized would require further investigation, but it is possible that, because the score distributions for essential and non-essential genes overlap, this gene was simply not hit enough times to achieve significance. This could be true for a small number of other genes as well and is a fundamental limitation of Tn-seq.

To further investigate our automated assignments, we screened eight “unassigned” genes in *S. islandicus* M.16.4 that were called essential by one method or the other but not both. We were unable to obtain mutants for six of them. Of these, five genes, i.e., *lig* (*M164_1953*), *priL* (*M164_1568*), *priX* (*M164_1652*), *rnhII* (*M164_0197*), and *tfs2* (*M164_1524*) were called essential via EI but not log_2_FC, while *thrS1* (*M164_0290*) was called essential based on log_2_FC but not EI. In contrast, knockouts of the two “unassigned” genes called essential by EI but not log_2_FC, *udg4* (*M164_0085*), encoding uracil-DNA glycosylase family 4, and *rpo8* (*M164_1872*), encoding a subunit of RNA polymerase, were obtained after an extended 14 days incubation of transformation plates, again consistent with a severe growth defect (Supplementary Fig. 2b, 2c, and 3b). This suggests the presence of false negatives and a stronger bias to underestimate than overestimate the true number of essential genes. Because not all genes in the unassigned categories were genetically tested, we conservatively excluded all unassigned genes from the essential gene list. By contrast, knockouts for all 76 non-essential genes tested were successfully obtained and verified by PCR analysis (Supplementary Table 3 and Supplementary Fig. 3). These include *hjm*/*hel308a* (*M164_0269*), *cdvB1* (*M164_1700*), *topR1* (*M164_1732*), and three DExD/H-box family helicase genes (*M164_0809*, *M164_2103*, and *M164_2020*), the homologs of which were previously thought to be essential in a related strain *S. islandicus* Rey15A^21–24^ (Supplementary Table 3 and Supplementary Fig. 3b). Taken together, these experimental results supported the overall validity of our computational approaches for conservatively classifying putative gene essentiality.

### Essential gene repertoire

The functional repertoire of the predicted essential, unassigned, and non-essential genes of *S. islandicus* is shown in Fig. 2. With the above adjustments, the size of this essential genome (441 genes) is close in size to that observed for other bacteria and archaea^25^. For example, ∼526 genes are required for growth in *Methanococcus maripaludis* S2^26^ and 473 genes within the engineered *Mycoplasma mycoides* JCVI Syn3.0 minimal bacterial cell^27^. The proportion of different functional categories represented in this set (as defined by archaeal clusters of orthologous genes^12^ (arCOG)) are also similar to that observed in other studies^26,27^ (Fig. 2), with the largest fraction of genes (178, 40%) representing information processing (translation, transcription, and DNA replication/recombination/repair) and 76 (∼17%) either classified as “Function unknown” or “General functional prediction only”. The latter two categories are hereby collectively referred to as “poorly characterized”. Descriptions of the specific essential components found in central information processing and the cell cycle, as well as central carbon metabolism, are detailed in Supplementary Information. We highlight only a few interesting and novel observations below.

**Figure 2:**
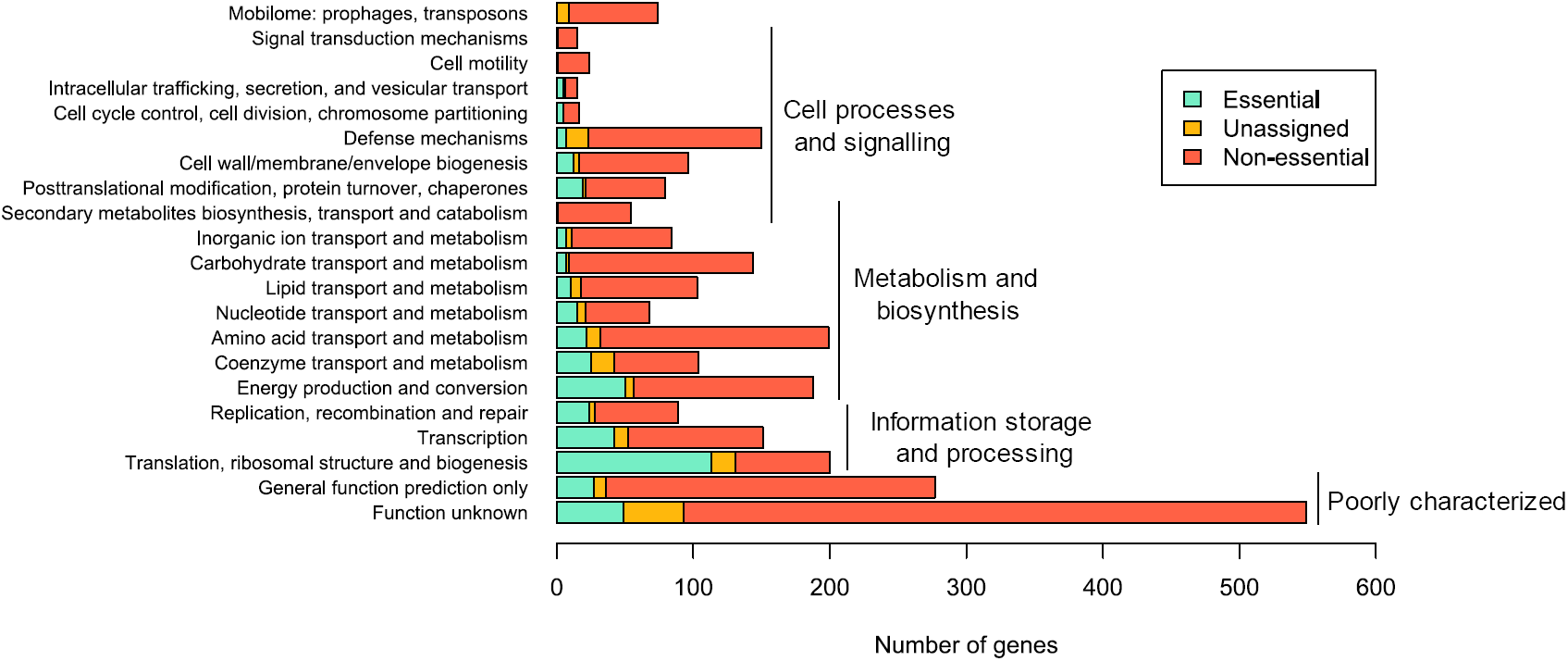
arCOG category and essentiality criteria for protein-coding genes in *S. islandicus* M.16.4. Functional distribution of essential, non-essential, and unassigned genes via arCOG category. Essentiality criteria based on cutoffs in Fig. 1b.

### S-layer is non-essential in *S. islandicus*

Our essential gene predictions include several surprising findings. First, SlaA (M164_1763) and SlaB (M164_1762), the two known components of the surface layer (S-layer) on the outside of *Sulfolobus* cells^28^ were shown to be non-essential. SlaA is the dominant component of the S-layer that forms a quasi-crystalline matrix the outside of the cell membrane^29^. Current models suggest a “stalk-and-cap” structure in which the C-terminal-transmembrane-helix-domain-containing SlaB projects from the cell membrane and anchors SlaA to the cell membrane^17,30,31^. The cellular function of the *Sulfolobus* S-layer is unknown, but is believed to provide resistance to osmotic stress and contribute to cell morphology^28^. S-layer deficient mutants have never been successfully cultivated before in any archaeal species, therefore it was assumed to be essential.

To confirm the non-essentiality of the S-layer genes, we constructed in-frame deletion mutants of *slaA*, *slaB*, and *slaAB* via a MID (<u>m</u>arker insertion and unmarked target gene <u>d</u>eletion) recombination strategy^32^. PCR amplification with two primer sets, which bind the flanking and internal region of S-layer genes, respectively (Fig. 3a), confirmed the successful deletion of *slaA*, *slaB*, and *slaAB* from the chromosome of the genetic host RJW004 (wild type) (Fig. 3b). We next tested for absence of the S-layer proteins in growing cells. Isolation of a white precipitant, described as the S-layer previously^33^, was possible only in the wild type and to a much lesser extent in the Δ*slaB* mutant strain (Supplementary Fig. 4a and 4b). Transmission electron microscopy (TEM) analysis confirmed this extracted protein precipitate from both wild type and Δ*slaB* formed crystalline lattice structures (Supplementary Fig. 4c). Finally, we tested the mutant phenotypes by comparing their growth profiles with wild type in a standard laboratory condition (pH 3.3, 76 °C). As shown in Fig. 3c, cells lacking the S-layer protein lattice SlaA (including *slaA* and *slaAB* mutants) are viable but have a measurable growth defect. This confirms the non-essentiality of the S-layer lattice in *S. islandicus*. The deletion of *slaB* alone had no significant impact on the growth rate in comparison with that of wild type (Fig. 3c). For a complete knockout of all potential S-layer components, we successfully created a viable triple knockout of *slaA*, *slaB*, and a paralog of SlaB encoded by *M164_1049* (42% coverage, 53% amino acid identity via BLAST), demonstrating non-essentiality of all S-layer components together in *S. islandicus* (Supplementary Fig. 5).

**Figure 3:**
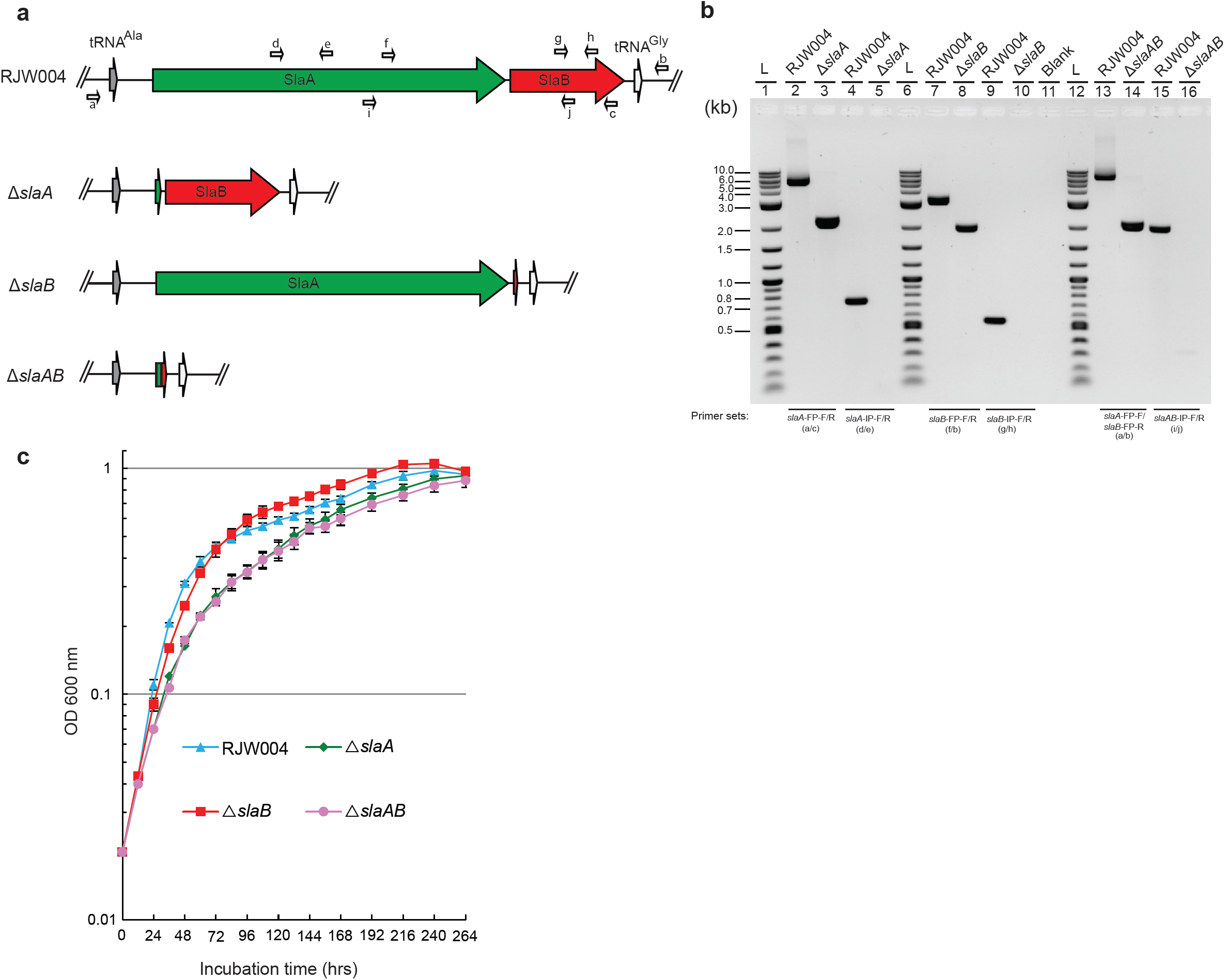
S-layer genes are not essential for the *S. islandicus* cell survivial. **a,** Genomic context of S-layer genes in the genetic host and mutant strains. Relative positions of primers used to confirm S-layer gene deletions are labelled with small arrows. **b,** PCR verification of Δ*slaA*, Δ*slaB*, and Δ*slaAB* mutants with two primer sets, which bind the flanking and internal regions of S-layer genes, respectively. Expected sizes of amplicons can be found in Supplementary Table 8. L (lanes 1, 6, and 12) indicates the 2-Log DNA Ladder (NEB, USA). Blank (lane 11) denotes that no sample was loaded in the well. **c,** Growth profiles of RJW004 (wild type), Δ*slaA*, Δ*slaB*, and Δ*slaAB* mutant strains. Wild type and S-layer gene knockout strains were cultivated at pH 3.3, 76 °C for 11 days in DY liquid medium supplemented with uracil and agmatine without shaking. Cell culture growth was monitored by optical density measurements at 600 nm every 12 or 24 hrs. Error bars represented standard deviations from three independent experiments.

We performed thin-section TEM analyses of the RJW004 (wild type) and S-layer gene knockout strains. The thin section micrographs of wild-type cells clearly revealed that the S- layer was separated from the cytoplasmic membrane by a quasi-periplasmic space (Fig. 4a and 4e), in agreement with previous studies in *Sulfolobus acidocaldarius*^34^ and *Sulfolobus shibatae*^29^. The S-layer in the wild type was observed as a distinct dark band on the outermost edge of the cell, and the quasi-periplasmic space was seen as a light grey band between the outermost band and the cell membrane (Fig. 4a and 4e). However, the dark, outermost layer surrounding the cell was not observed in the Δ*slaA* or Δ*slaAB* mutant cells (Fig. 4b, 4d, 4f, and 4h), confirming that SlaA contributes to the formation of the outermost layer. Additionally, the cell surface appeared diffuse in the Δ*slaA* mutant cell, which was attributed to the periodic extensions of membrane proteins, likely including the SlaB protein, and/or their extensive N- glycosylation^17^. In the Δ*slaB* mutant cell, a smooth outermost layer similar to the SlaA layer in wild-type cells was observed; however, it appears to be discontinuous around the cell membrane (Fig. 4c and 4g). The partial lattice of SlaA in the Δ*slaB* mutant may be anchored by other membrane proteins even in the absence of SlaB, including the aforementioned M164_1049. Together these images suggest additional components may contribute to the non-essential *Sulfolobus* S-layer^17^.

**Figure 4:**
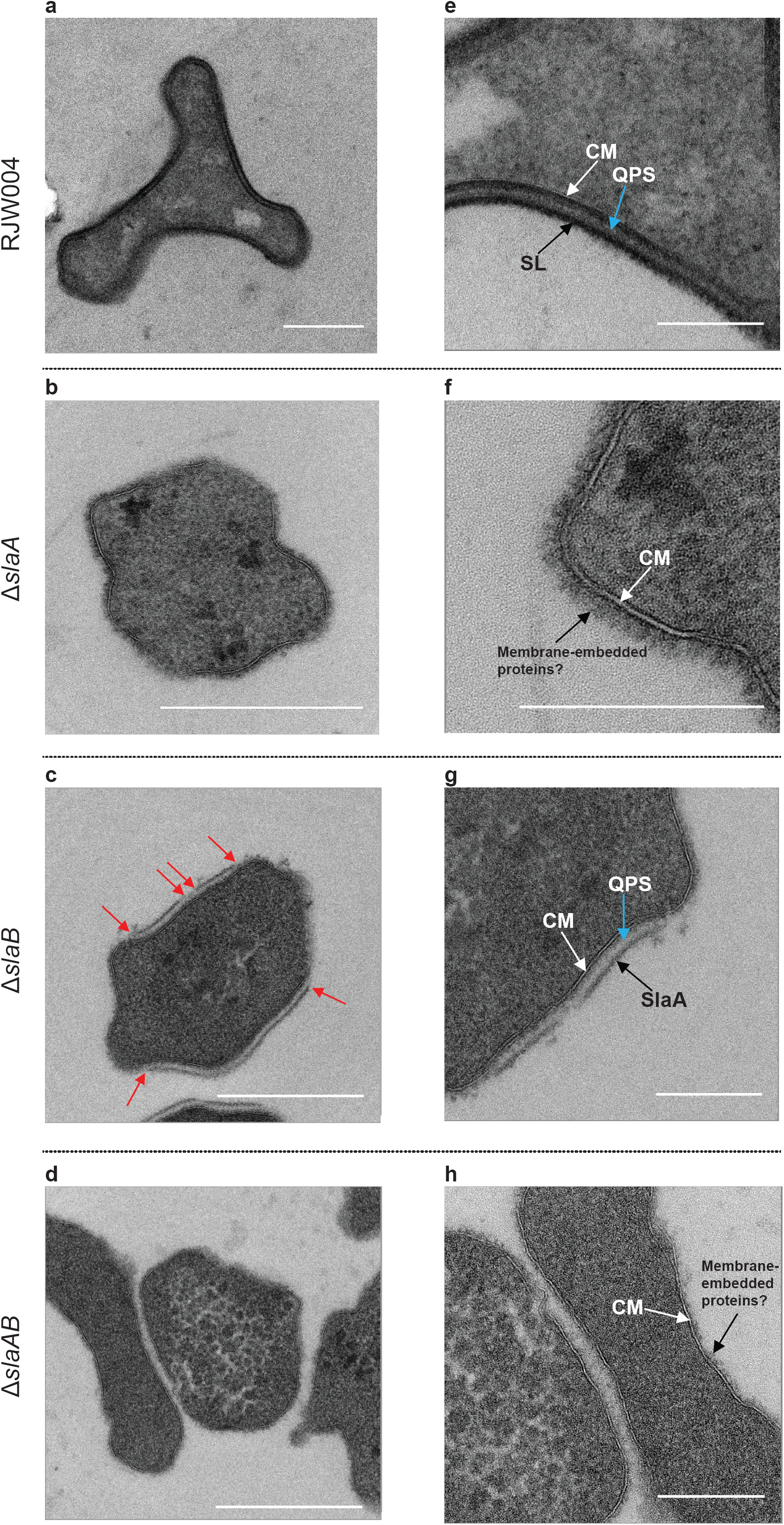
Thin-section TEM analysis of the wild type and S-layer gene knockout strains. **(a-d),** Representative TEM micrographs of thin-sectioned cells of the wild type, Δ*slaA*, Δ*slaB*, and Δ*slaAB* mutant strains, respectively. Images **(e-h)** are closeups of images **(a-d)**, respectively. Red arrows indicate the breaking points of S-layer. Abbreviations: CM, cytoplasmic membrane. SL, surface layer. QPS, quasi-periplasmic space. SlaA, surface layer protein A. Scale bars, 500 nm **(a-d)**, and 200 nm **(e-h)**.

### Incomplete complementarity of reverse gyrase

As an additional surprise from our genome-wide essential gene identification, we found incomplete complementarity between two copies of the reverse gyrase in *S. islandicus* M.16.4. Unlike Euryarchaota and most extremely thermophilic bacteria, Crenarchaeota possess two copies of reverse gyrase^35, 36^, both believed to be essential for growth^21, 37^. Tn-seq analysis indicated that the *topR1* (*M164_1732*) was non-essential, which was confirmed by a successful disruption (Supplementary Fig. 1b). Interestingly, as mentioned above, *topR2* (*M164_1245*) was called essential but we could obtain *topR2* disruption mutants (Supplementary Fig. 1c) if we prolonged the incubation time (up to 14-20 days) of transformation plates in gene knockout experiments. These observations suggest that *topR2* plays a more important role than *topR1* in *Sulfolobus* cell survival at optimal temperature.

### Lethal deletion mutants

Tn-seq also uncovered genes that may not be essential to growth but instead are toxic when disrupted. Among them, arCOG analysis predicts that *M164_0131*, *M164_0217*, *M164_0268*, *M164_2076*, *M164_1728*, and *M164_1060* are antitoxin-encoding genes. We reason that inactivation of these antitoxin genes might cause overproduction of toxins and then trigger cell death. This finding suggests that associated toxins are constitutively expressed in our laboratory conditions. Interestingly, unlike most of the family II (VapBC) and family HEPN-NT toxin/antitoxin family gene pairs in *S. islandicus* M.16.4 (Supplementary Dataset 3), partners (toxin genes) adjacent to these predicted antitoxin genes (with the exception of *M164_1060*; see Supplementary Fig. 6) were not observed. This indicates that VapB-VapC or HEPN-NT do not always correspond to their neighbors and some gene pairs might have exchanged counterparts. The Tn-seq-based analyses also classified *cas5* (*M164_0911*), a part of the Cascade (CRISPR-associated complex for antiviral defense) complex^38^, as essential. Consistent with this assignment, disruption of *cas5* by replacing it with the *StoargD* marker cassette via homologous recombination failed after repeated attempts. However, the entire Type-IA module of CRISPR-Cas system, consisting of eight genes with *cas5* included, could be deleted from the *S. islandicus* M.16.4 chromosome with no detectable effect on cell growth (data not shown). One possible explanation is that in the absence of *cas5*, the Cascade complex becomes misfolded and thus toxic for the cells, but future studies are needed to confirm this interpretation.

### Shared essential genes

To establish how this essential gene set compares with those found in other organisms, we retrieved sets of essential genes from the database of essential genes^25, 39^ in 8 model organisms that span the tree of life^26,40–44^, including the minimal genes set in the JCVI Syn 3.0 *Mycoplasma mycoides* genome^27^ (Fig. 5, Supplementary Dataset 4). We find that 242 *S. islandicus* essential genes are essential in at least one other organism we surveyed, while 192 essential genes are uniquely essential in *S. islandicus*. Eighty-nine genes are essential in representatives of all three domains (78 of which are also essential in Syn 3.0) (Supplementary Dataset 4). As shown in Fig. 5, comparisons of shared essential genes support the shared cellular systems between the archaeal and eukaryotic domains. More total *S. islandicus* essential gene orthologs are shared with archaea and eukaryotes (grey in Fig. 5), and more of these shared orthologs are essential (colors), than are shared between *S. islandicus* and the bacteria we use for comparison. The highest number of shared essential genes (187) is between *S. islandicus* and *M. maripaludis* S2^26^, an organism from the euryarchaeal lineage of the archaeal domain (Table 1). The large size of the essential gene set shared between *Sulfolobus* and *Methanococcus*, in spite of their wildly different habitats and life styles, reinforces the fundamental nature of Archaea as a distinct cell type^4^.

**Figure 5:**
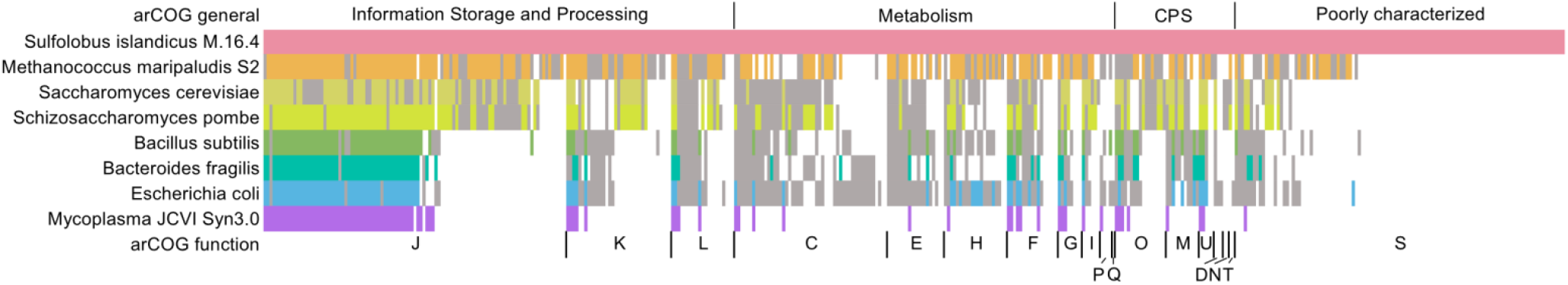
Shared essential genes across the three domains of life. Heatmap shows the presence of essential (colored) or non-essential (grey) shared NOGs compared with the *S. islandicus* essential genome. Single-letter codes for functional categories are as follows: J, translation, ribosomal structure and biogenesis; K, transcription; L, DNA replication, recombination, and repair; C, energy production and conversion; E, amino acid transport and metabolism; H, coenzyme transport and metabolism; F, nucleotide transport and metabolism; G, carbohydrate transport and metabolism; I, lipid transport and metabolism; P, inorganic ion transport and metabolism; Q, secondary metabolites biosynthesis, transport and catabolism; O, post-translational modification, protein turnover, chaperone functions; M, cell wall/membrane/envelope biogenesis; U, intracellular trafficking, secretion, and vesicular transport; D, cell cycle control and mitosis; N, cell motility; T, signal transduction; S, function unknown; CPS, cellular processes and signaling.

**Table 1:** Number of *S. islandicus* essential genes shared and shared essential within 8 model organisms.

|  |  |  | Phyletic Category* |  |  |  |  |  |
| --- | --- | --- | --- | --- | --- | --- | --- | --- |
|  |  |  | Universal | EA | Archaea | TACK | Sulfolobales | Other |
| Archaea | *Methanococcus maripaludis* | Shared | 128 | 77 | 42 | 2 | 2 | 42 |
|  |  | Essential | 93 | 55 | 20 | 2 | 0 | 17 |
| Eukarya | *Saccharomyces cerevisiae* | Shared | 134 | 77 | 2 | 0 | 1 | 27 |
|  |  | Essential | 68 | 41 | 1 | 0 | 0 | 4 |
|  | *Schizosaccharomyces pombe* | Shared | 134 | 78 | 2 | 0 | 1 | 26 |
|  |  | Essential | 82 | 43 | 1 | 0 | 0 | 10 |
| Bacteria | *Bacteroides fragilis* | Shared | 124 | 10 | 0 | 0 | 2 | 40 |
|  |  | Essential | 70 | 1 | 0 | 0 | 0 | 12 |
|  | *Bacillus subtilis* | Shared | 131 | 7 | 5 | 1 | 4 | 49 |
|  |  | Essential | 72 | 0 | 0 | 0 | 0 | 11 |
|  | *Escherichia coli* | Shared | 136 | 7 | 6 | 0 | 8 | 64 |
|  |  | Essential | 73 | 0 | 1 | 0 | 0 | 12 |
|  | *JCVI Syn 3.0* | Shared | 76 | 2 | 0 | 0 | 0 | 10 |
|  |  | Essential | 76 | 2 | 0 | 0 | 0 | 10 |

\* Genes are put into a category if they are present in >50% of the organisms in each group, i.e. universal is in >50 % of each of Bacteria, Archaea and Eukarya groups. "Other" refers to genes that do not meet these criteria.

### Phyletic distributions of essential genes

To investigate the broader phyletic distributions of *S. islandicus* essential genes, we used assignments from the eggNOG database^46^ (see Methods) to map the presence and absence of putative essential gene orthologs from a previously published set of 169 complete genomes representing major clades in all three domains^6^ (Supplementary Dataset 5 and Supplementary Dataset 6). Fig. 6 graphically shows the *S. islandicus* essential genes shared in other genomes in a set of hierarchical clusters based on Euclidean distance. From this figure, 4 primary transitions emerge in the evolution of the contemporary S. *islandicus* essential genome. The number of genes in phyletic groups (Table 2) is significantly different from random sampling among phyletic categories (Supplementary Table 4). Similarly ranked distributions are seen in two additional datasets: 1) all genomes in the eggNOG database subsampled to have equal representation in each domain, 2) all genomes in the eggNOG database for which assignments are available. These data are supported by parsimony analysis with bootstrap support for the grouping of each of the three major domains (Supplementary Fig. 7 and 8). Together these data support four primary stages in the evolution of the contemporary *S. islandicus* cells and allow us to assign specific essential genes to these potential transitions in the evolution of the cell.

**Figure 6:**
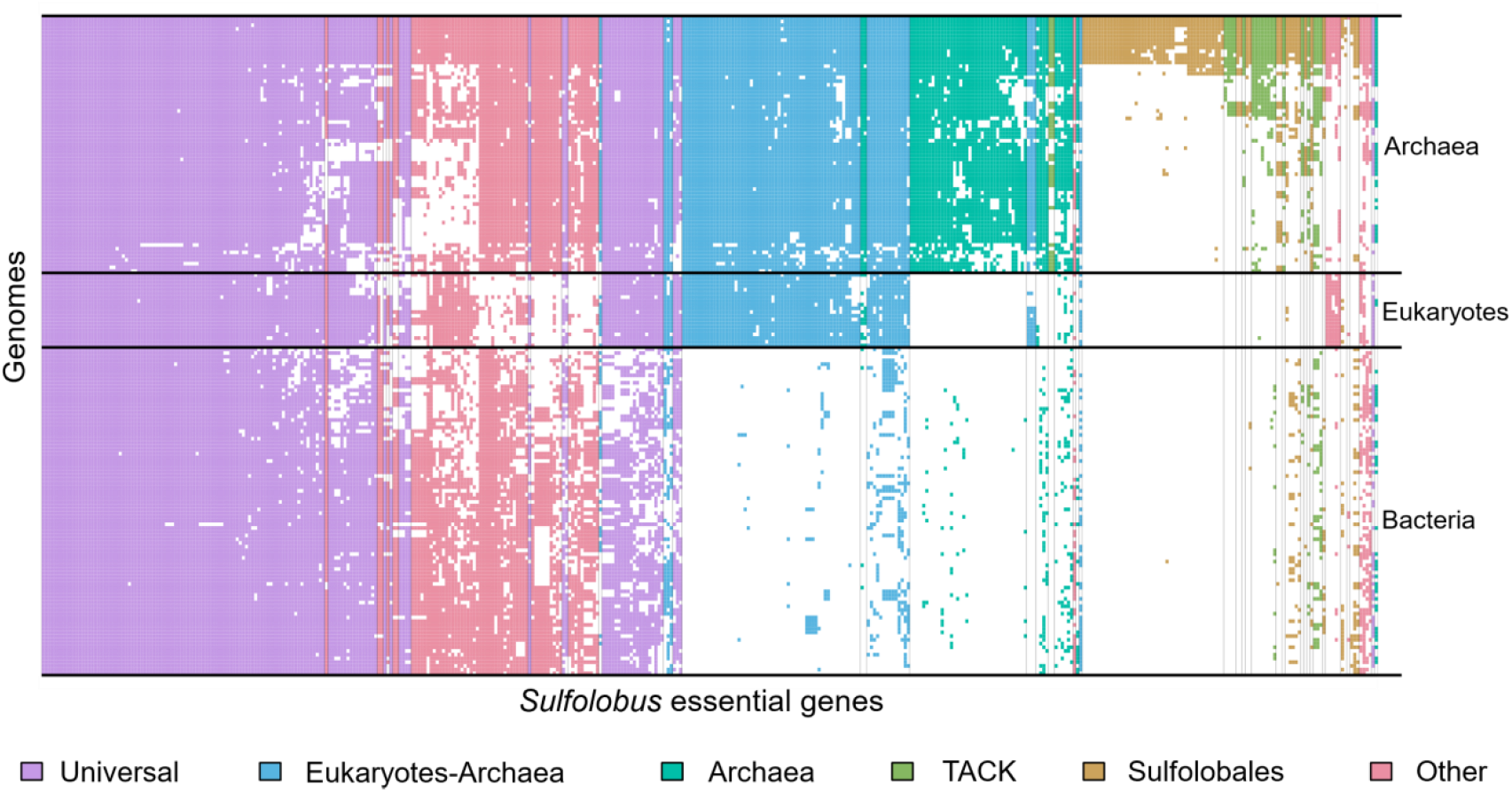
Presence/absence of genes shows phyletic patterns. Heatmap of shared NOG/arNOGs according to annotations in the eggNOG database corresponding to the essential gene set of *S. islandicus* across the three domains of life. Each row is one of 177 taxa including the set of 169 used for other distribution analyses, one from the candidate phylum *Bathyarchaeota*, and 7 Asgardarchaeota genomes (Supplemental Datasets 5 and 6). Each column is one of 441 essential genes discovered in this study. A white box indicates no matching NOG or arNOG was found, while a colored box indicates presence. Colors indicate categories defined in Table 1 and Table 2.

**Table 2:** Number of *S. islandicus* essential genes shared with 168 full genome sequences spanning tree of life.

|  | Phyletic Category* |  |  |  |  |  |
| --- | --- | --- | --- | --- | --- | --- |
|  | Universal | EA | Archaea | TACK | Sulfolobales | Other |
| **Share Genes** | **141** | **80** | **55** | **18** | **73** | **74** |
| **Poorly Characterized\*** | 5 | 1 | 14 | 7 | 46 | 3 |

\* Genes are put into a category if they are present in >50% of the organisms in each group, i.e. universal is in >50 % of each of Bacteria, Archaea and Eukarya groups. "Other" refers to genes that do not meet these criteria.
\*\* NOG categories "Function unknown" or "General functional prediction only". Full list shown in Supplemental Table 6

The highest number of essential genes are shared broadly across the tree of life (Universal in Table 2), supporting the early evolution of the majority of essential gene functions in the contemporary archaeal cell. Most of these have putative functional assignments in information processing, particularly translation and transcription (Supplementary Dataset 7). Many previous studies have reported the evolutionary conservation of information processing components going back to the Last Universal Common Ancestor (LUCA) using computational methods^7–9,11,47^. We find that in all studies the majority of conserved orthologous gene sets that we could interrogate in this system are essential (Supplementary Table 5 and Supplementary Dataset 8). Of the 200 metabolic COGs identified in the *S. islandicus* genome from a recent estimate of the LUCA gene set^10^, only 19 were found to be essential (Supplementary Dataset 8). This is expected, due to our use of rich medium. The first phase of the cell contains the universal set of genes with conserved cellular components that are likely to have evolved early in evolutionary history remain essential components of the contemporary *S. islandicus* genome today.

The next largest category of essential genes is found between *Sulfolobus* and other organisms in the Eukarya/Archaea (EA) domains (Table 2). These genes are largely involved in core information processing functions and support the shared evolutionary ancestry of the Archaea and Eukarya after their divergence from Bacteria. Only one gene in this category is poorly characterized: *M164_0237*, a homolog to eukaryotic *zpr1*. *zpr1* is a gene essential for transcription and cell cycle progression in fungal and mammalian cells^48-51^, and has recently been reported as a regulator of circadian rhythm in plants^52^. Though it has been noted that this gene is exclusively shared in EA^51^, it remains uncharacterized in the Archaea outside of our results recognizing its essentiality in *Sulfolobus* (Supplementary Table 6).

Fifty-five essential genes belong to NOGs that are shared by organisms in the archaeal domain (Table 2). Functional assignments of the archaeal-specific genes represent a diversity of functions split between core functions (translation, transcription, and replication) and peripheral functions such as transport, defense (including all the above-mentioned predicted antitoxin genes), and metabolism. Archaea-specific DNA replication/recombination/repair genes are *nurA* and *gins15*, while genes in arCOG category “Transcription (K)” are largely transcription factors and do not represent core RNA polymerase functionality like the EA genes mentioned above. Fourteen of the archaeal-specific genes are poorly characterized (Table 2), 9 of which are also essential in *M. maripaludis* S2 (Supplementary Dataset 4). In an evolutionary context, this set of poorly characterized, but essential, archaea-specific genes are key target for future molecular characterization since they likely highlight the unique biology of archaeal cells. We also show that the majority of *S. islandicus* genes are conserved in evolutionary history through the archaeal domain.

The final set of essential genes are specific or largely specific to the Sulfolobales, most of which have uncharacterized functions (Table 2). The essentiality of these genes and whether they fit into central cellular functions as non-orthologous gene replacements or peripheral ones are important subjects of future work. This set of genes, unique to this lineage, may represent environmental adaptations. The fact that they are poorly characterized attests to the need for further study even in this model archaeon.

The phyletic distributions of essential gene orthologs describe more about the shared biology of organisms than about the evolutionary processes (invention, loss, and horizontal gene transfer) through which each combination of essential components evolved. The key next steps toward comparative cell biology will be understanding the functional interactions among essential genes so that new gene inventions, non-orthologous gene transfers, and/or loss of specific functions can be identified. From the unique perspective of the TACK archaea, this work provides a roadmap of genes whose future molecular and systems characterization are likely to provide further understanding for evolutionary steps in the Archaea.

## Conclusion

This is the first comprehensive genome-wide study of essential gene content in a model crenarchaeon. Our profile of *S. islandicus* essential genes uncovers several surprising findings, most notably the non-essentiality of the *Sulfolobus* S-layer. Comparative phyletic patterns provide a perspective on the stages of evolution of the contemporary *S*. *islandicus*, its shared ancestry with the eukaryotes, and the key components that define its uniqueness as an archaeal cell.

## Methods

### Strains and culture conditions

The complete list of strains and plasmids used in this study is shown in Supplementary Table 7. All *S. islandicus* strains were routinely grown aerobically at 76-78 °C and pH 3.3 without shaking in basal salt medium^32^ containing 0.2% [wt/vol] dextrin (Sigma-Aldrich, USA) and 0.1% [wt/vol] tryptone (BD Biosciences, USA) (the medium is hereafter named as DY). When required, agmatine, uracil, and 5-FOA were added to a final concentration of 50 μg/ml, 20 μg/ml, and 50 μg/ml respectively. For solid plates, 2 × DY medium was supplemented with 20 mM MgSO_4_ and 7 mM CaCl_2_·2H_2_O, and mixed with 1.4% gelrite (Sigma-Aldrich, USA) with a ratio of 1:1 [vol/vol]. Plates were put into sealed bags and generally incubated for 10-14 days at 76-78 °C. Cell culture growth was monitored by optical density measurements at 600 nm using a portable cell density meter (CO8000, WPA, Cambridge, United Kingdom).

### Methods

#### Construction of *S. islandicus* transposon mutant library

The 755-bp *argD* gene cassette (*SsoargD*) was PCR-amplified from the genomic DNA of *S. solfataricus* P2 using the primer set *SsoargD*-F1/R1, introducing the SalI and XmalI sites respectively. The resultant PCR products were digested with SalI/Xmal, and then cloned into the EZ-Tn5^TM^ pMOD^TM^-2 <MCS> Transposon Construction Vector (Epicentre, USA) in the corresponding sites, generating pT-SsoargD. The Tn5 <SsoargD> transposon DNA was prepared by PCR amplification from linearized pT-SsoargD with 5’phosphorylated primers PCRFP/PCRRP. The PCR products, consisting of a nutritional marker flanked by a 19-bp inverted repeat (Mosaic Ends, ME), were purified and highly concentrated using the DNA Clean & Concentrator^TM^-5 kit (Zymo Research, USA). Preparation of transposomes was made in a 10 μl-reaction system as follows: 2.2 μg of transposon, 1 μl of EZ-Tn5 transposase (Epicentre, USA), and 2.5 μl of 100% glycerol. The reaction was incubated at room temperature for 30 min and then switched to 4 °C for another 72 hrs. 1-2 μl of transposomes were transformed into *S. islandicus* RJW008 (Δ*argD*) via electroporation as described previously^32^. Cell transformation assays were repeated dozens of times in order to collect a sufficient number of transformants to achieve saturation mutagenesis. The theoretical number of transposon insertion colonies was calculated using a derivative of Poisson’s law: N = ln(1 - P)/ln(1 - f), f = average gene size (900.64 bp) / genome size (2,586,647 bp). To make sure the transposon insertions cover approximately 99.99% (P=0.9999) of the genome, around 26,448 colonies per library are required. The transformed cells were plated on DY plates either by glass beads or over-lay^53^. After 10 days of incubation, the ArgD^+^ revertants were harvested from plates either by manually picking or with sterile spreaders, and then pooled into three independent transposon mutant libraries (CYZ-TL1, CYZ-TL2, and CYZ-TL3), with approximately 100,000 colonies in total (Supplementary Table 1). We routinely obtained an average of *ca.*10^3^ colonies/μg transposon, and approximately 10^5^ colonies/μg DNA using a replicative plasmid pSeSd-SsoargD. The pSeSd-SsoargD was constructed by cloning the *SsoargD* marker cassette, amplified from *S. solfataricus* P2 genomic DNA with primer set *SsoargD*-F2/R2, into the XmalI site of a *Sulfolobus-E.coli* shutter vector pSeSd^54^. Thus, the estimated frequency of transposition is ∼10^-2^ per cell.

#### DNA library preparation and high-throughput DNA sequencing

Genomic DNA from each mutant pool was extracted as described previously^55^ and then quantified with Qubit^®^ 2.0 Fluorometer (Invitrogen, USA). DNA libraries were prepared using the Nextera XT DNA Library Prep Kit (Illumina, USA) with proper modifications. Briefly, 2 ng of input genomic DNA in total was simultaneously fragmented and tagged with sequencing adapters in a single enzymatic reaction tube. Afterwards, a primer mixture of Tn-seq-F (Supplementary Dataset 9) and N705 (a randomly selected primer from the Nextera XT DNA Library Prep Kit) was added in the same tube to enrich the transposon-chromosome junction regions via PCR. The PCR conditions were as follows: 72°C for 3 minutes, 95°C for 30 seconds, and 22 cycles of denaturation at 95°C for 10 seconds, annealing at 55°C for 30 seconds, and extension at 72°C for 30 seconds. A final extension was performed at 72 °C for 5 min. The resultant library DNA was cleaned up with AMPure XP beads for three times, eluted in 45-μl EB buffer (QIAprep Spin Miniprep Kit, USA), and then quantified with Qubit^®^ 2.0 Fluorometer. The final DNA library was quantitated on High-Sensitivity Qubit (Life Technologies) and fragment size was evaluated using the Agilent 2100 Bioanalyzer on a DNA7500 chip (Agilent Technologies), then further quantitated by qPCR on a BioRad CFX Connect Real-Time System (Bio-Rad Laboratories, Inc. CA) to ensure accuracy of quantitation of the library containing properly adapted fragments. The final pool was loaded onto two lanes (CYZ_TL1) and 1 lane each (CYZ_TL2 and CYZ_TL3) of a HiSeq 2500 Rapid flowcell for cluster formation and sequencing on an Illumina HiSeq 2500 with Rapid SBS sequencing reagents version 2. Sequencing by synthesis was performed from one end of the molecules for a total read length of 160 nt. The 100 μM of custom Read 1 sequencing primer, specific for the Tn-seq-F sequence (Supplementary Dataset 9), was spiked into the standard Read1 HP10 primer tube (position 18) for sequencing. The run generated .bcl files, which were converted into demultiplexed compressed fastq files using bcl2fastq v1.8.4 Conversion Software (Illumina, CA) at the W. M. Keck Center for Comparative and Functional Genomics at the University of Illinois at Urbana-Champaign.

#### Tn-seq data processing and analysis

Illumina FASTQ reads from all three libraries that were fewer than 50 bp in length, had a quality score below 30, and did not contain the 23-bp transposon sequence were removed. The remaining reads were stripped of transposon and adapter sequence and aligned to the *S. islandicus* M.16.4 genome (NC_012726) using the Burrows-Wheeler Bowtie 2 alignment tool^56^. Reads that mapped to multiple locations in the genome or to ambiguous sites were set aside, as were those with an alignment length less than 11 base pairs. Using in-house software, the resulting .sam alignment files were converted to lists that included unique insertion locations, the strand to which they aligned, and the number of reads associated with that event (Supplementary Dataset 1). Insertions that occurred in the same location but on different strands or in separate libraries were considered independent events. Tn5 transposase has been shown to prefer certain insertion sites over others^57^, so each reported site was extracted and nucleotide frequency was measured 20 bases up-and-downstream as compared to an equal number of random sites in the genome. Random sampling via the Python numpy.random.choice function (with replacement) yielded sites with overall frequencies matching the known G+C content of the genome (35%), but a pronounced and palindromic pattern was observed at insertion sites even when normalizing for this bias (Supplementary Fig. 9). Overall Tn5 appears to prefer a G-C base pair flanked by an AT-rich region, which is consistent with other studies^57, 58^. However, when normalized to the overall G+C content, no single biased site was more than 2-fold enriched in a certain base compared to the rest of the genome, meaning there was considerable variation in the sites themselves and thus the chance that the bias would significantly affect our results is reduced. Gene essentiality was then evaluated using software previously designed and published for this purpose: Tn-Seq Explorer^19^ and ESSENTIALS^18^. The ESSENTIALS software was run with mostly default settings with a list of insertion locations and associated reads as the input. The locations for each of the three libraries were submitted as separate files and the total library size specified as 105,968 (Supplementary Table 1). Repeat filtering was enabled to avoid calling repeated regions as essential. The LOESS smoothing feature normally meant to compensate for the over-representation of bacterial origins of replication (caused by multiple simultaneous replication rounds) was disabled because *Sulfolobus* only undergoes one round of replication per cell cycle^59^. Because of the lack of observed sequence specificity, the insertion site was specified as “random.” The program uses “log_2_FC” as its measure of essentiality, which is proportional to log_2_ (reads observed/reads expected) for each gene and sets a cutoff automatically as the local minimum between essential and non-essential distributions in a density plot of the scores. The program suggested a putative maximum log_2_FC of -5.1 for essential genes.

For the Tn-Seq Explorer software, insertion sites of all three libraries were combined and insertion sites with fewer than 4 reads were excluded for analysis due to their vast over-representation in the insertion sites and the uncertainty of their source (Supplementary Table 1). The program uses a sliding window approach and returns an essentiality index (EI) based on the number, location, and spatial concentration of insertion sites within each individual gene. It also allows for the adjustment of the stated start and end points of the gene. As is default, insertions in the first 5% and last 20% of genes were excluded to compensate for misannotated start codons and proteins for which C-terminal deletions are tolerated, respectively. The program suggested an EI maximum of 3 (Fig. 1b).

#### Construction of gene replacement and markerless in-frame deletion mutants in *S. islandicus*

Except where otherwise states, disruption of the chromosomal genes was achieved by replacing their coding regions (57%-100% of the length of the gene was deleted) with the *argD* expression cassette (*StoargD*) derived from *S. tokodaii* via a microhomology-mediated gene inactivation approach we recently developed^60^. Briefly, a functional *argD* gene was PCR-amplified from a linearized *Sulfolobus*-*E.coli* shuttle vector pSesD-StoargD with 35-40 bp homology of the targeted gene introduced, yielding the gene disruption cassettes. The resultant PCR products were purified and electroporated into the *argD* auxotrophic strain *S. islandicus* RJW008, selecting ArgD^+^ transformants on the plates lacking agmatine. S-layer genes *slaA*, *slaB*, and *slaAB* were deleted from the chromosome of the genetic host *S. islandicus* RJW004 via an improved MID strategy^32, 61^ with knockout plasmids pMID-slaA, pMID-slaB, and pMID-slaAB, respectively. The resulting Δ*slaA* and Δ*slaB* mutants harbored an in-frame deletion of the coding region from nucleotides +52 to +3687 relative to the start codon of *slaA* (3690 bp in length), and +13 to +1185 relative to the start codon of *slaB* (1194 bp in length), respectively. The Δ*slaAB* mutant was constructed similarly leaving 51 bp of the *slaA* (nt 1 to 51 relative to the start codon of *slaA*), 6 bp of restriction enzyme (MluI) site, and 9 bp of *slaB* (nt 1186 to 1194 relative to the start codon of *slaB*) in the chromosome of *S. islandicus* RJW004. Verification of each gene replacement or deletion mutant was determined through PCR diagnosis with both flanking primers (bind outside of the targeted region) and internal primers (bind inside of targeted region), which examined the genotype and purity of mutants respectively. The primers used to generate and confirm gene disruptions or deletions were described in the Supplementary Dataset 9, and the expected sizes of amplicons generated from the genetic host (wt) and mutant strains were provided in the Supplementary Table 8.

#### Transmission electron microscopy (TEM)

Proteinaceous S-layer was extracted from *S. islandicus* cell cultures as described previously^33^. To prepare the samples that were used for TEM, glow-discharged, carbon-stabilized Formvar-coated 200-mesh copper grids (Carbon Type-B, cat. no. 01811, Ted Pella, Inc., USA) were placed on 8- to 20-μl droplets of each sample for 3 minutes, rinsed with deionized water, and negative-stained with 2% uranyl acetate for 15-60 seconds. Thin-sectioned *S. islandicus* cells were prepared essentially as described previously^62^, with minor modifications as follows: after microwave fixation with the primary fixative, cells were washed in Sorenson’s Phosphate buffer with no further additives. All samples were observed using a Philips CM200 transmission electron microscope at 120 kV. Images were taken at various magnifications using a TVIPS (Tietz Video and Image Processing Systems GmbH; Germany) 2k × 2k Peltier-cooled CCD camera. Scale bars were added with ImageJ software.

#### Homology search

Homologs for the 441 essential genes found in Supplementary Dataset 2 were found across the 168 genomes listed in Supplementary Dataset 6 via the European Molecular Biology Laboratory evolutionary genealogy of genes Non-supervised Orthologous Groups (EMBL eggNOG) database^46^. Genomes were downloaded from the National Center for Biotechnology Information (www.ncbi.nlm.nih.gov). The genomes to survey are based on the set^6^ in Raymann*, et al.* 2015 with the following additions: *M. maripaludis* S2 was added to compare essential gene content; the genome of *Toxoplasma gondii* ME49 was added because its essential genome became available during the course of this analysis^63^; *Schizosaccharomyces pombe*^64^ was added to compare with *Saccharomyces cerevisiae* S288C; additional Sulfolobales genomes were added for intra-order comparison of essential gene content (listed in Supplementary Dataset 6). While not included in the phyletic distribution analysis, the sequences for *Lokiarchaeum* sp. GC14_75^14^ and Thorarchaeota^65^ SMTZ-45, SMTZ1-45, and SMTZ1-83 were retrieved and analyzed; presence/absence data can be found in Supplementary Dataset 5. Several bacterial genomes were added to include additional model systems (e.g. *E. coli* str. K-12 substr. MG1655 and *Bacillus subtilis* subsp. *subtilis* str. 168). The complete list is found in Supplementary Dataset 6. Due to their incomplete or highly reduced nature, we excluded DPANN and Asgard lineages, as well as Bacteria from the candidate phyla radiation^66^ and the minimal *Mycoplasma* Syn 3.0^27^; however, presence/absence information for selected genomes are provided in Supplementary Dataset 5. For organisms not in the eggNOG database, the amino acid sequences of protein-coding genes were uploaded to the eggNOG mapper tool (http://eggnogdb.embl.de/#/app/emapper) and run with default settings. These data were translated into a presence/absence matrix and evaluated with custom Python and Zsh scripts to assess the phyletic distribution of essential gene candidates. Finally, for each *S. islandicus* M.16.4 essential gene candidate, the amino acid sequences of all bidirectional best BLAST hits of that gene were used to scan genomes in which no homologs were found search using tBLASTn, and the results were filtered according to the same cutoff criteria as the bidirectional best BLAST hits. This was to fill in gaps left by annotation mistakes, where the protein may still be in the genome but was not published as such. The tBLASTn hits that overlapped with annotated genes by more than 50 base-pairs were discarded because the search was explicitly for finding missing annotations. Presence/absence patterns of NOG homologs were combined with tBLASTn data to create binary matrices.

#### Parsimony analysis

Presence/absence matrices were converted to NEXUS format files with a custom Python script and used in the Phylogenetic Analysis Using Parsimony (and other methods) (PAUP*) tool^67^. The main tree was found with the heuristic search function with a maximum of 1000 trees in memory, default settings. The first tree was saved as an unrooted NEXUS format tree with branch lengths. Bootstrapping was run with default settings for 1000 iterations with 100 maximum trees in memory. The resulting tree was saved with support values as node labels. A custom python script using the Phylo package within the Biopython^68^ suite was used to transfer the support values from the bootstrap consensus tree to the corresponding nodes on the heuristic search tree. Trees were visualized in the interactive tree of life (iTOL)^69^ interface.

#### Phyletic distribution analysis

The presence/absence matrices were also cross-referenced with phylogenetic data via NCBI taxonomy information to determine how widespread each gene was in different orders spanning the tree of life. Simulated random distributions of genes were created by counting how many organisms in which they were found and assigning that many random organisms to each gene (without replacement) for each gene 100 times using the numpy.random.choice function. P-values were generated by counting the number of simulated observations above or below the true observation and dividing by 100. To determine the proportion of COGs or arCOGs that have essential members in *S. islandicus*, we removed unique clusters with more than one gene in *S. islandicus* that showed no essentiality from the total due to possible redundancy in functional orthologs from the same cluster. To test for bias in the phyletic sampling of the set of 169 genomes, we assembled 100 organism sets with genomes randomly sampled from the eggNOG database to equal proportions of TACK archaea, euryarchaeota, eukaryotes, and bacteria as to that in the 169 genome set. Organisms were chosen at random without replacement using the numpy.random.choice function again. In all data involving the eggNOG database, organisms missing from the NOG members file were excluded.

## Life Sciences Reporting Summary

Further information on experimental design is available in the Life Sciences Reporting Summary.

## Code availability

The custom Python and Zsh scripts used for analyses in this study are available upon request.

## Data availability

The raw Tn-seq data of three independent transposon insertion libraries CYZ-TL1, CYZ-TL2, and CYZ-TL3 have been deposited at NCBI under BioSample accessions SAMN08628694, SAMN08628695, SAMN08628696, respectively, Bioproject accession PRJNA436600, and Sequence Read Archive (SRA) accession SRP133799. Analyzed data showing the insertion locations across three independent transposon libraries can be found in Supplementary Dataset 1. All other data that support the findings of this work are available from the corresponding author upon request.

## Acknowledgments

We thank Chris L. Wright and Alvaro G. Hernandez from W.M. Keck Center for Comparative and Functional Genomics, University of Illinois at Urbana-Champaign (UIUC), for advice with primer design, DNA library construction and sequencing. We also thank Whitney E. England, Angelo Blancaf and Ted Kim for assistance in Tn-seq data processing; Carlos A. Vega, Elizabeth H. Marr, and Melinda E. Baughman for providing technical assistance in collecting transposon insertion colonies. We thank Isaac K.O. Cann and Scott C. Dawson for fruitful discussions. We are thankful to Kira S. Makarova for technical assistance with data retrieval from the arCOG database and for helpful suggestions. We thank Marleen van Wolferen for providing the S-layer extraction protocol. We acknowledge Yuan Li and Emily N. Hallett for providing S-layer extraction assistance. We would also like to thank Scott J. Robinson from Beckman Institute for Advance Science and Technology, UIUC, for technical assistance with TEM imaging and sample preparation. We thank Lou A. Miller from Frederick Seitz Materials Research Laboratory Central Research Facilities, UIUC, for preparing the thin-sectioned *S. islandicus* cells. Funding for this work was mainly provided by the National Aeronautics and Space Administration (NASA) through the NASA Astrobiology Institute under cooperative agreement no. NNA13AA91A, issued through the Science Mission Directorate. This work was also partially supported by Division of Environmental Biology (DEB: 1355171 to R.J.W.), US National Science Foundation, the Department of Microbiology Alice Helm Graduate Research Excellence Fellowship, UIUC (to A.P.R.P.), the Carl R. Woese Institute for Genomic Biology Undergraduate Research Scholar program, and the Office of Undergraduate Research, UIUC (to R.L.W).

## Author Contributions

C.Z., A.P.R.P., and R.J.W. conceived and designed the research; C.Z. and R.L.W. carried out experimental work; A.P.R.P., C.Z., G.J.O., R.L.W., and R.J.W. analyzed the data; R.J.W. and G.J.O contributed new reagents/analytic tools; and C.Z., A.P.R.P., and R.J.W. wrote the paper. All authors edited the manuscript.

